# Actin dynamics as a multiscale integrator of cellular guidance cues

**DOI:** 10.1101/2021.05.26.445782

**Authors:** Abby L. Bull, Leonard Campanello, Matt J. Hourwitz, Qixin Yang, Min Zhao, John T. Fourkas, Wolfgang Losert

## Abstract

Cells are able to integrate multiple, and potentially competing, cues to determine a migration direction. For instance, in wound healing, cells follow chemical signals or electric fields to reach the wound edge, regardless of any local guidance cues. To investigate this integration of guidance cues, we monitor the actin-polymerization dynamics of immune cells in response to cues on a subcellular scale (nanotopography) and on the cellular scale (electric fields, EFs). In the fast, amoeboid-type migration, commonly observed in immune cells, actin polymerization at the cell’s leading edge is the driver of motion. The excitable systems character of actin polymerization leads to self-propagating, two-dimensional wavefronts that enable persistent cell motion. We show that EFs guide these wavefronts, leading to turning of cells when the direction of the EF changes. When nanoridges promote one-dimensional (1D) waves of actin polymerization that move along the ridges (esotaxis), EF guidance along that direction is amplified. 1D actin waves cannot turn or change direction, so cells respond to a change in EF direction by generating new 1D actin waves. At the cellular scale, the emergent response is a turning of the cell. For nanoridges perpendicular to the direction of the EF, the 1D actin waves are guided by the nanotopography, but both the average location of new actin waves and the whole cell motion are guided by the EF. Thus, actin waves respond to each cue on its intrinsic length scale, allowing cells to exhibit versatile responses to the physical microenvironment.

**Significance Statement:** Effective cell migration requires the integration of multiple, and sometimes competing, guidance cues. For instance, in wound healing, immune cells are guided towards a wound edge by long-range electrical and chemical cues that may conflict with guidance cues from the local environment. How cells combine and respond to such cues is not well understood. We demonstrate that multiple guidance mechanisms can act simultaneously, but on different scales. Nanotopography, a local mechanical cue, guides individual waves of actin polymerization, thereby biasing the direction cell motion on the time scale of these waves. An external electric field applied at the same time biases the locations of new waves of actin polymerization, leading to overall directed migration over long distance scales.

## Introduction

Cells often operate as part of a collective organism, in which individual cells carefully coordinate their behaviors with each other and their environment. This coordination between cells and their environment is modulated not only by well-known biochemical signals, but also through a range of additional cues, including the physical properties of the local environment. One important, highly regulated cell behavior is migration, which can be guided by cues such as chemotactic signals (1), electric fields (EFs) (2), and the texture of the local environment (3). Recent studies of cell migration in developing *Drosophila* have demonstrated that, *in vivo*, cells can find a path of least resistance in an environment with adhesive, topographic and chemoattractant signals (4). Here we investigate how cells integrate information when multiple guidance cues are present simultaneously. We focus on neutrophils, which naturally experience several guidance cues simultaneously (chemical signals, EFs, and the texture of the extracellular matrix) when migrating toward a wound. The guidance induced by the chemical signals and EF produced by a wound is typically in the same direction. However, local guidance cues, such as from the extracellular matrix, do not necessarily point in the same direction. To develop a deeper understanding of this competition, we focus on the manner in which neutrophils integrate and respond to the combination of a local guidance cue, nanotopography, and a guidance cue that operates over a longer distance, an external EF.

The signaling pathways that drive cell motility converge in actin polymerization, enabling this process to integrate multiple migratory cues. Recent research has demonstrated that these signaling pathways include both fast, positive feedback loops, and slow, negative feedback loops (5). These signaling pathways are coupled to the cytoskeleton, driving cytoskeletal rearrangements that enable cells to perform tasks such as developing integrin-mediated adhesions, forming protrusions, elongating, polarizing intracellular components, and migrating (6– 9). Recent studies have demonstrated that signaling pathways and the actin polymerization machinery together act as an excitable system, with characteristic waves and oscillations of actin polymerization (10)(11). The hallmarks of excitability are an all-or-none response to an external signal and a refractory period after excitation. Such behavior is characteristic of actin polymerization, which occurs in waves and oscillations on a time scale of tens of seconds and over a distance scale of microns.

In excitable signaling networks, upregulation of a single protein can be sufficient to change the characteristic length scale of actin-polymerization waves. Such a change in wave scale can in turn alter the migratory phenotype dramatically, from local filopodia driven by sub-micron scale actin polymerization, to lamellipodia on the order of several microns, to protrusions spanning the entire cellular periphery (7, 12). Because small changes in the balance of signaling molecules can be sufficient for a cell to change from one migratory phenotype to another, measurements of actin-wave characteristics offer a comprehensive view into the state of a cell that can be more directly related to cellular physiology than quantification of, for instance, protein concentrations.

Here we study the manner in which actin waves respond to combinations of EFs and nanotopography. In wound healing, EFs with magnitudes in the range of 0.5 to 2 V/cm can prompt directed cell migration via electrotaxis (13). EFs may further activate signaling pathways that lead to changes in the polarization of cellular components (14, 15). Prior work has offered conflicting opinions regarding the importance of actin polymerization in the cellular sensing of EFs (16, 17). EFs have been found induce cell polarity by localizing the polarity indicator PI3K (PIP_3_) toward the cathode in neutrophil-like HL60 cells, even when the cells are treated with latrunculin, an inhibitor of actin polymerization (2). In both untreated and latrunculin-treated HL-60 cells, the response to an EF direction reversal was observed to have an induction time of several minutes. However, latrunculin-treated *Dictyostelium discoideum* cells were found not to localize polarity markers towards the cathode (18). These observations indicate that actin polymerization plays a direct role in the response to EFs under at least some circumstances.

Cells are also known to be guided by the second guidance cue employed here, nanotopography. As opposed to cell behavior on flat surfaces, migration guided by one-dimensional topographies yields cell morphologies and cytoskeletal organization that closely mimics guided migration in a three-dimensional environment (19). In neutrophils, the response to nanotopography enables effective migration to wound sites through complex extracellular microenvironments, including collagen fibers and the extracellular matrix (3, 20). *In vivo*, cells do not simply follow guidance provided by nanotopography, they can also reshape it. For example, neutrophils have been observed to rearrange collagen networks in wound environments to allow for wound closure (3).

Here we use parallel nanoridges with dimensions comparable to typical extracellular matrix fibers (21) to guide local actin dynamics, in a process called esotaxis. Esotaxis promotes the formation of guided, one-dimensional waves of actin polymerization that are distinct from the two-dimensional waves observed on flat substrates, and that are therefore likely more representative of the actin dynamics *in vivo*. Esotaxis leads to waves with widths on the submicron scale of the individual topographical features, enhancing the directionality and persistence of cell migration for a wide range of cell types (22–26).

We investigate actin dynamics when EFs and nanoridges are combined either in a cooperative manner (parallel to one other) or a competitive manner (perpendicular to one another). We demonstrate how the excitable-systems character of the cell migration machinery allows EFs to provide guidance over long temporal and spatial scales even in the presence of strong local guidance cues from nanotopography. These results may help to explain the ability of the electric fields of wounds to promote healing even in the presence of competing local guidance cues.

## Results

We measured the response of the actin dynamics of neutrophil-like HL60 cells, an established model for migratory immune cells (27), to EFs and nanotopography. The nanoridges had widths comparable to those of collagen fibers (300 nm width, spaced 1.5 μm apart), and the EFs were comparable in magnitude to those found in healing wounds. We exposed actin-YFP neutrophils to a 10 V/cm EF that first pointed from right to left (anode to cathode) for 5 min (Fig. 1*Ai*). We reversed the EF direction for the remaining 10 min of the experiment. After EF reversal, the actin cytoskeleton rearranged the location of the cell front over several minutes, which caused the cell to turn and migrate from left to right (Fig 1*Aii-iv*). Kymographs of a region along the EF axis (yellow boxes in Fig. 1*A*) show slanted lines of high intensity at the leading edge of the cell that reappear in the opposite direction a few minutes after the EF reversal (Fig. 1*A* right, green arrows). On nanoridges with a 1.5 µm spacing aligned parallel to the EF (Fig. 1*B*), the actin dynamics is less coordinated, and the cell lacks a well-defined and connected actin front (Fig. 1*Bi*). In particular, the uniform actin polymerization along the front of the cell that was visible initially (first green arrow, Fig. 1*B* right) appears to break up into localized, one-dimensional actin waves during the turn, consistent with the quasi-one-dimensional waves seen on nanoridges in a broad range of cell types (25). These individual actin waves can respond to the EF independently of one another (Fig. 1*Bii, iii*). Although individual actin waves only travel along the axis of the nanoridges, the cell overall undergoes a turning motion. After several minutes, a uniform actin wavefront reemerges, oriented towards the new cathode (Fig. 1*Biv*).

**Figure 1.**
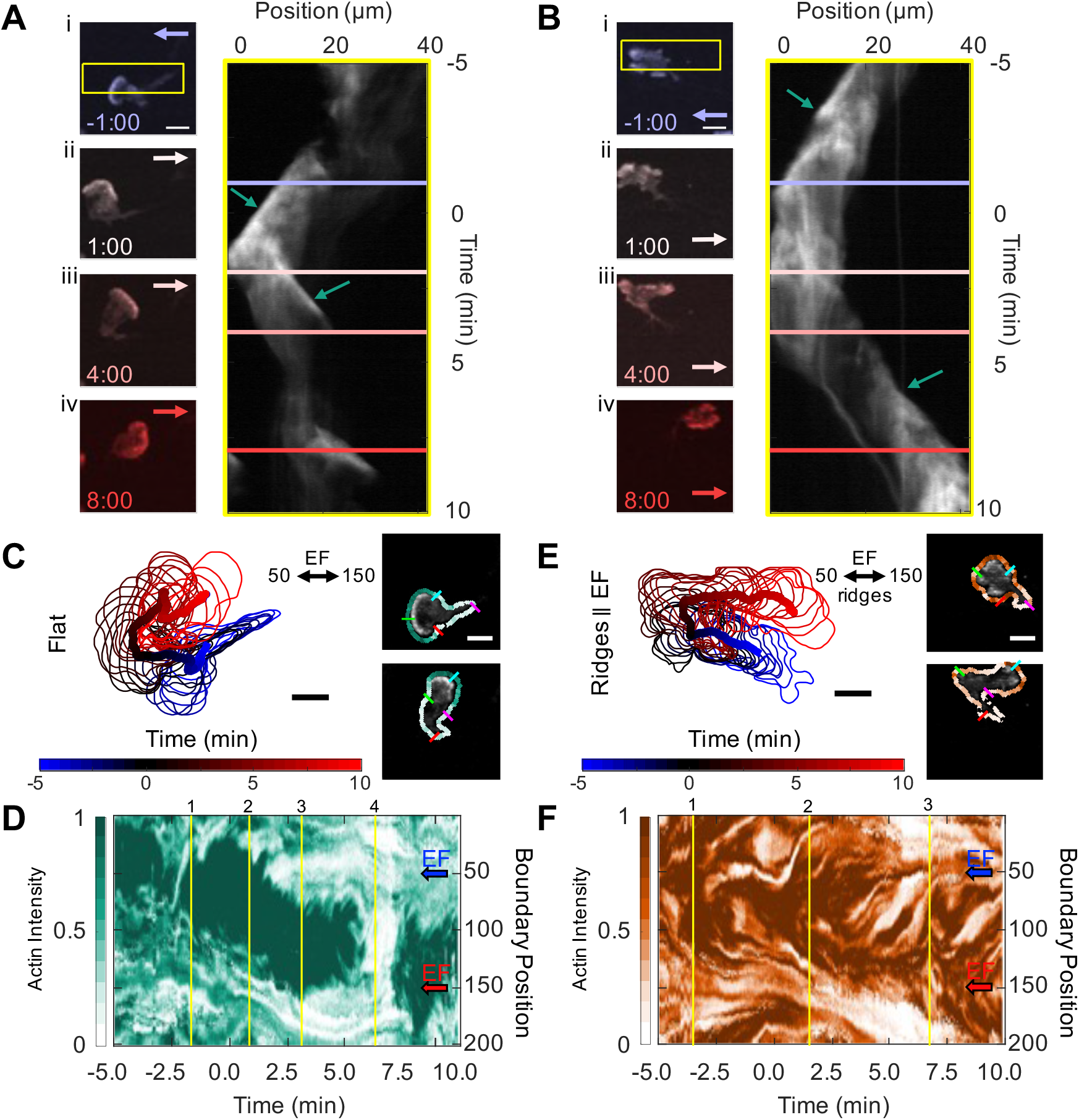
HL60 cells turn toward the new cathode within several minutes of EF reversal on both flat and ridged substrates, but with different local actin structures. (A) Four time-lapse snapshots of a differentiated, YFP-actin-labelled HL60 cell on a flat substrate. The cathodal (negative) direction is to the left at negative times. The EF reversal occurred at time zero. Kymograph (right) for area denoted by the yellow border in (i). (B) A differentiated, YFP-actin-labelled HL60 cell on nanoridges parallel to the EF. Kymograph (right) for area denoted by the yellow border in (i). Scale bars are 10 µm. The arrows in (A) and (B) denote the EF direction, and the corresponding time slices in the actin intensity kymographs. (C) Boundary shape and path of the centroid of cell from (A) from -5:00 min to 10:00 min in 0.5-min increments, with an example of the extracted shape of a cell with labeled indices of boundary at points 50, 100, 150 and 200 on right; the boundary is colored by actin intensity. (D) Evolution of normalized boundary actin fluorescence intensity from (A,C) visualized as a kymograph with notable features shown with yellow lines. Lines 1-3 show the shifting actin intensity around the cell and line 4 highlights a cringe period of the cell. (E) Boundary shape and path of the centroid of cell from (B) from -5:00 min to 10:00 min in 0.5-min increments with an example of an extracted cell shape with labeled indices of boundary points 50, 100, 150 and 200 on right; the boundary is colored by actin intensity. (F) Evolution of normalized boundary actin fluorescence intensity from (B,E) visualized as a kymograph. Line 1 shows splitting and merging of actin waves before the EF switch. Line 2 shows a wave in the wrong direction (top) that disappears by time 5.0 and a lower wave that continues to propagate as the cell orients to the cathode. Line 3 indicates the low-actin rear of the cell that appears after the EF reversal. The blue and red arrows in (D, F) indicate the cathodal direction at the specified boundary points for negative time and positive time, respectively. Scale bars: 10 µm.

Active-contour analysis of the evolving shape of the cells provides additional insights into the differences in response to changing EF directions by enabling visualization of the turning of a cell (Fig. 1*C*) (28, 29). To link the evolving shape of a cell with actin dynamics, the actin concentration surrounding each point along the cell membrane was measured at each time step (Fig. 1*C* right). The actin dynamics associated with the turning cell can then be visualized through a kymograph of averaged actin intensity along the cell boundary (Fig. 1*D*). As the cell turns, the boundary region with the highest actin concentration shifts in a clockwise fashion (times 1–3, yellow lines). This continuous turn appears as a diagonal band in the kymograph. The cell then undergoes a rounding-up, or cringe, period (time 4, yellow line), with a uniform, low level of actin fluorescence.

The evolving shape of the cell on nanoridges from Fig. 1*B* exhibits a similar turning motion (Fig. 1*E*), but with centroid motion that is altered by the nanotopography. Following the reversal of the EF, the centroid of the cell undergoes rapid transitions from moving to the left, to moving perpendicular to the ridges, to moving to the right. During the turn, actin near the cell boundary is distributed in multiple patches (Fig. 1*E* right). The time dependence of the actin fluorescence intensity around the boundary of the cell on nanoridges (Fig. 1*F*) indicates that these actin waves split and merge even before the EF direction is changed (time 1, yellow line). During the turning of the cell, these multiple actin waves do not appear to turn, but rather compete with one another. Waves oriented in the anodal direction disappear after several minutes (line 2, top dark region), and waves oriented in the cathodal direction persist (line 2, lower dark region). Line 3 indicates the reappearance of the low-actin-concentration (i.e., light-colored), contractile tail of the cell, completing the EF-driven reorientation of the cell. In addition, for cells that are rounded, we have observed behavior consistent with activation of neutrophils, i.e., a transition from rounded to polarized morphology with the initial introduction of an EF (Fig. S1).

For a more thorough analysis of the differences in actin dynamics on flat and ridged substrates, it is useful to quantify the spatial and temporal intracellular changes in an unbiased manner. For this purpose, we employed optical-flow analysis (25, 30, 31), as shown in Fig. 2. Using spatiotemporal actin fluorescence intensity gradients as inputs (Fig. 2*A*), optical-flow fields enable quantitative characterization of the actin-wave dynamics (Fig. 2*B*). The distribution of directions of the actin dynamics is shown in Fig. 2*C* for three representative cells in 5-min increments. The analysis shows a change in preferred direction of actin dynamics that follows the changing cathode direction. Fig. 2*C*(left) shows the actin dynamics for the cell in Fig. 1*A*. The actin flow changes its direction of motion from the left to the upper right, with an intermediate turning period in which the flow is oriented roughly equally left and right. In Fig. 2*C* (middle), the actin flow changes from left to right, with the flow at the intermediate time interval oriented less accurately with the EF. In Fig. 2*C* (right), the cell undergoes a left-to-right flip in the flow between the first- and second-time intervals (blue to black), after which the flow loses its left/right bias and is oriented perpendicularly to the EF. Conversely, on nanoridges, the flows orient more accurately along the ridge/EF axis. In Fig. 2*D* (left), the flow distribution for the cell in Fig. 1*B* is quantified in the same three time periods. Compared to the flat surface, there is stronger guidance of the actin waves in the direction of the EF on nanoridges. In Fig. 2*D* (middle), the flow is initially bidirectional, and then shifts to the right in the third time interval. In Fig. 2*D* (right), the flow is guided unidirectionally by the EF before the field reversal (blue). After the field switch, the flow continues in the leftward direction (black), before becoming bidirectional along the ridges in the third time interval (red). Although the actin dynamics is polarized in each cell, the direction of polarity is different for each cell on flat surfaces. Thus, on a flat surface, the cumulative results for all cells show only a slight EF guidance of actin waves (Fig. 2*E*). In contrast, on nanoridges that is a strong bidirectional guidance of actin dynamics with a clear preference for flow toward the cathode (Fig. 2*F*).

**Figure 2.**
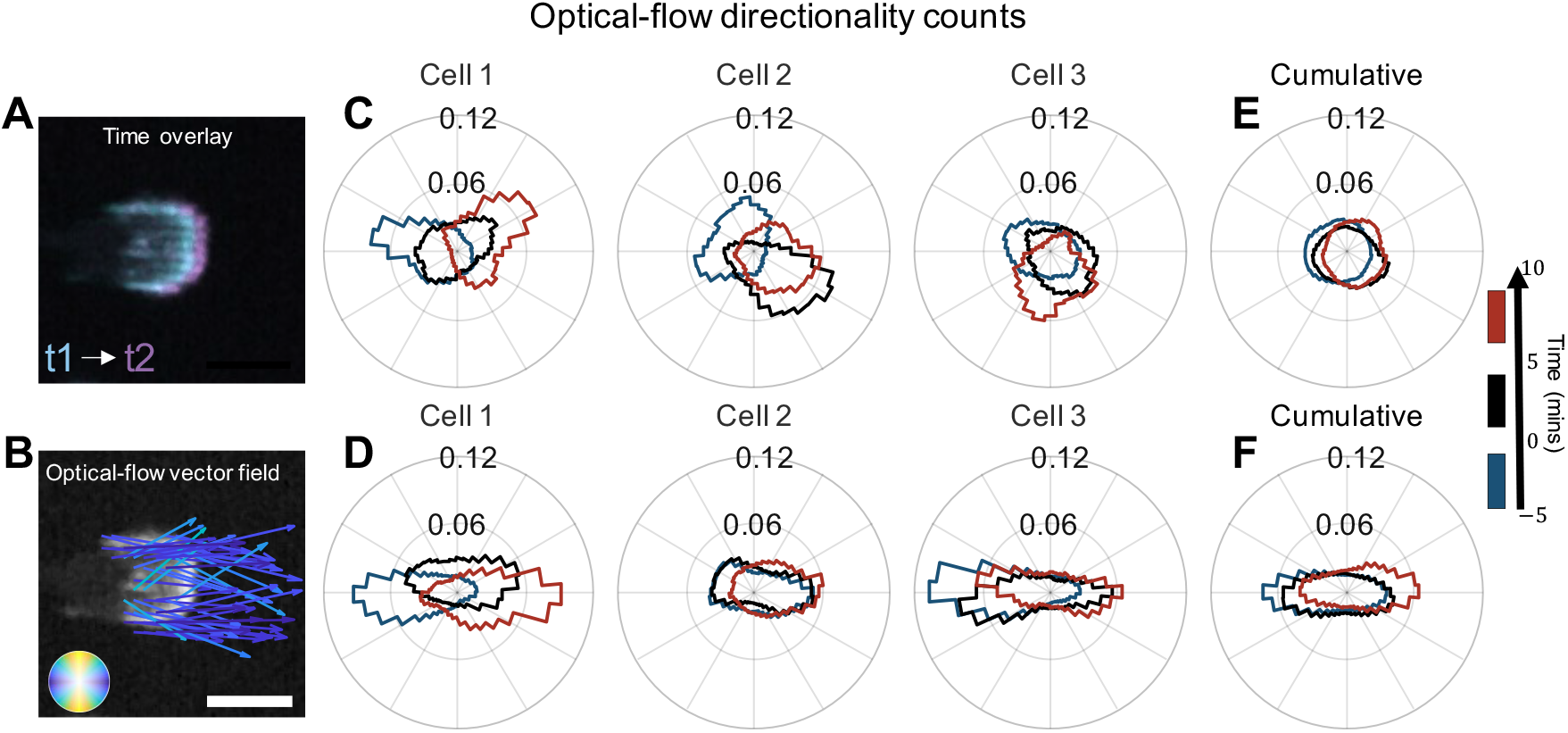
Nanotopography aligned with an EF enhances guidance of actin dynamics. (A) Time overlay of two actin-fluorescence images of an HL60 cell on nanoridges; the temporal spacing is 0.5 min. (B) Representative optical-flow vectors for the motion between the frames in (A). Scale bars: 10 µm. Three characteristic examples of individual, cell-normalized distributions (in counts) of optical-flow directions on a flat surface (C) and nanoridges (D) (with the leftmost histograms from cells in Fig. 1) shown for the time ranges: (blue) 5 min before the EF reversal; (black) 5 min after the EF reversal; and (red) the period from 5 min to 10 min after the EF reversal. The cumulative flow-direction distribution for the same time intervals for all cells on (E) the flat surface (n = 15) and (F) the nanoridges (n = 11).

We investigated how actin dynamics respond to two competing guidance cues by orienting the nanoridges perpendicular to the EF. Fig. 3*A* shows actin fluorescence images of a representative cell before and after EF reversal. Before EF reversal, the cell is oriented with its front towards the cathode (Fig. 3*Ai*). After EF reversal, new actin waves emerge on the side of the cell facing the new cathode (Fig. 3*Aii*). Once the previous actin-rich regions disappear (Fig. 3*Aiii*), a wide region of actin appears across the leading edge of the cell, facing in the new cathode direction (Fig. 3*Aiv*). The trajectory of the cell boundary and centroid of the cell are shown in Fig. 3B. The kymograph of cell-boundary actin intensity in Fig. 3*C* displays a jump in actin intensity on the boundary from left (50) to right (150) shortly after the EF reversal. Although the location of actin patches is on the side of the cell closest to the cathode, the nanoridges strongly affect the dynamics of the patches. Actin-flow distributions for three representative cells (Fig. 3D) and cumulative flow distributions (Fig. 3*E*) show strong alignment of the actin flow with nanoridges, with a slight preference for flow in the direction of the EF. See Fig. S2 for similar quantification for cells responding to the initial application of the EF.

**Figure 3.**
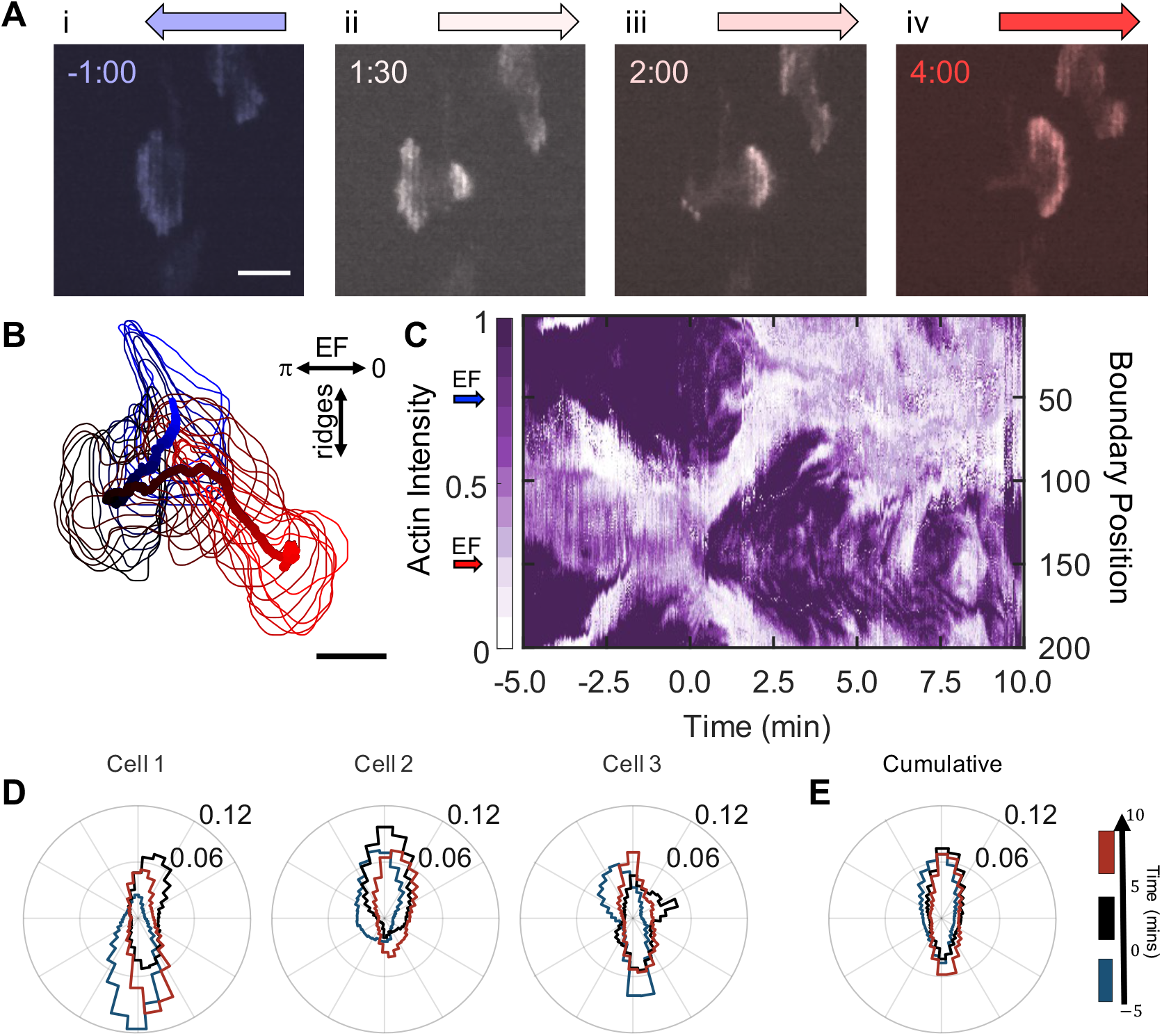
Nanoridges dominate the local guidance of actin waves in the presence of competing EF guidance. (A) A YFP-actin-labelled, differentiated HL60 cell on nanoridges aligned perpendicularly to the EF. The cathodal direction is to the left at negative times; the EF reversal occurred at time zero. (B) Path of a cell centroid with boundary shape from -5 min to 10 min, plotted every 0.5 min. (C) Evolution of normalized boundary actin fluorescence intensity. (D) Three characteristic examples of individual cell normalized distributions (in counts) of optical-flow directions on perpendicular ridges, shown for the time ranges: (blue) 5 min before the field was reversed; (black) 5 min after the field was reversed; and (red) the period 5 min to 10 min after the field was reversed – leftmost distribution for cell in (A)-(C). (E) The cumulative flow-direction distribution for the same time intervals for all cells (n = 9).

To gain further insight into the competition between the guidance of the cell by the EF and of the actin by nanoridges, we reanalyzed the actin dynamics with a larger smoothing length scale for all three combinations of guidance cues (EF on flat substrates, EF parallel to nanoridges, and EF perpendicular to nanoridges). For the actin-flow measurements in Figs. 2 and 3, we initially smoothed the dynamics over a scale of 0.6 µm, which is smaller than the ridge spacing of 1.5 µm (Fig. 4*A*). For that optical-flow calculation, the cumulative flow directions for all three combinations of guidance cues indicate guidance of actin flow by the nanoridges, but only limited guidance by an EF perpendicular to the nanoridges (Fig. 4*B*). However, when the actin dynamics is smoothed on a 2.1-µm scale, which is greater than one ridge spacing, some guidance by a perpendicular EF becomes apparent (Fig. 4*C* and supplemental Fig. S3). To capture actin dynamics on the length scale of single cells, we tracked the centroid of actin intensity over time. For this tracking, we excluded dim actin in the cell bulk to focus on the brighter regions at the forefront of motion. In this case, guidance by both nanoridges and EF orientation is notable when these guidance cues compete (Fig. 4*F*).

**Figure 4.**
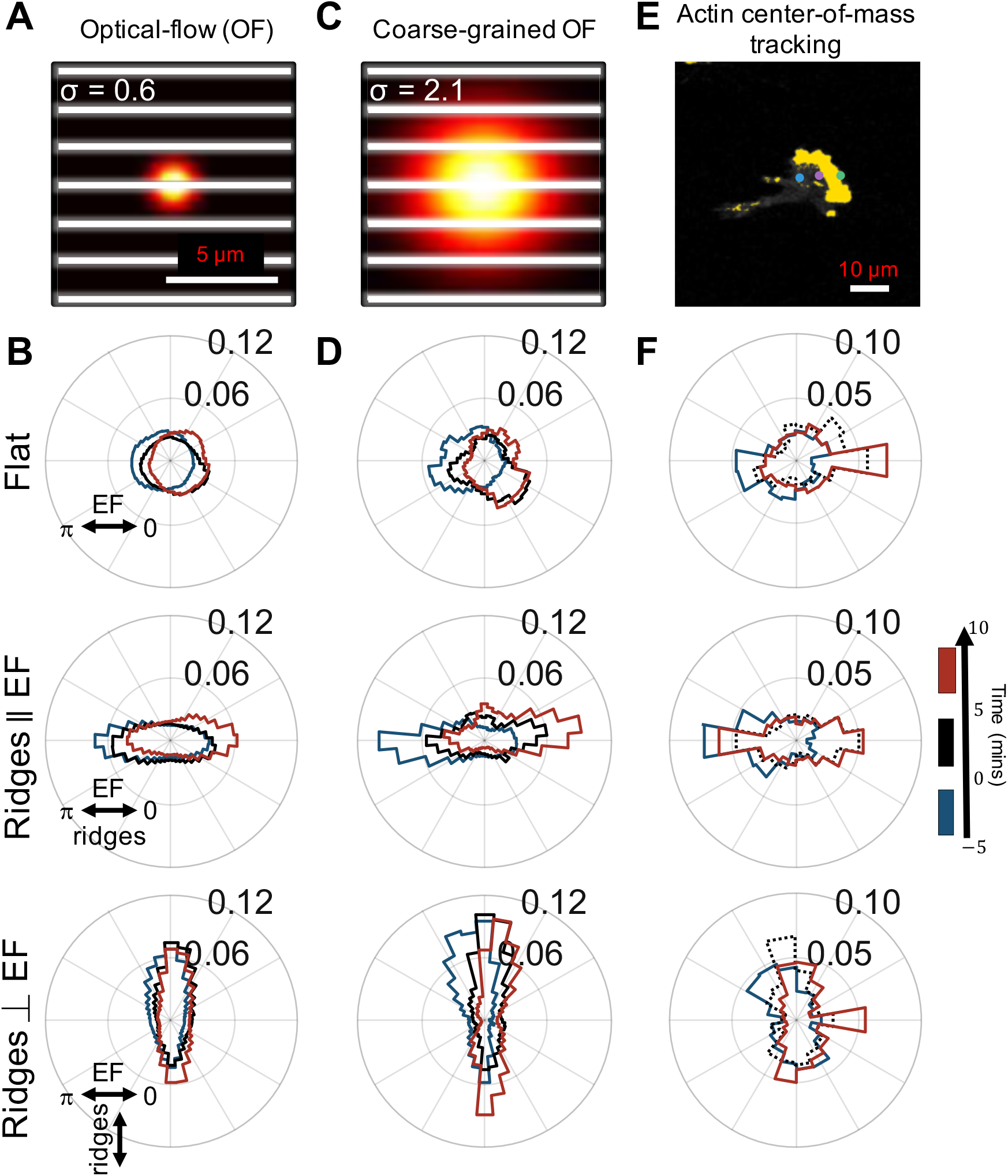
Actin dynamics analyzed on multiple scales with cooperating and competing nanotopography and EF guidance. Actin polymerization is predominantly guided by nanotopography, but the guidance is biased by the EF. (A) Visualization of spatial scale used for optical-flow calculations (white lines = ridges). (B) Cumulative flow-direction distribution shown for the three-time intervals, as previously described for all cells on (top) a flat substrate, (middle) nanoridges parallel to the EF, and (bottom) nanoridges perpendicular to the EF. (C) Visualization of the spatial scale used for coarse-grained optical-flow calculations to capture larger-scale actin waves. (D) Cumulative flow-direction distribution shown for the same time ranges as (B) for all substrate conditions. (E) Visualization of actin center-of-mass tracking via the Crocker-Grier algorithm (39), with track results from three time points shown. (F) Actin centroid tracking for all substrate conditions.

To highlight the guidance by the EF direction in the presence of significant cell-to-cell variability, we calculated the logarithm of the ratio of leftward and rightward dynamics of actin, which we denote *R*. This metric highlights the existence of a response to the EF reversal, and does not depend on the accuracy of the response. As above, we compare the actin dynamics in the time period of 5 to 10 min after the EF reversal to those in the 5 min immediately prior to the reversal. A scatter plot of *R* for optical-flow measurements smoothed on the 0.6-µm scale (OF scale) versus *R* for the same cell on the cellular scale (actin COM) highlights the significant cell-to-cell variability, and at the same time reveals that dynamics on both scales are correlated for all three guidance conditions (Fig. 5*A*). The strongest correlation across spatial scales is observed for nanoridges oriented parallel to the EF. Note that some cells do not follow the directional guidance of the EF. Specifically, *R* is smaller than zero for blue triangles and larger than zero for red circles. The strong correlation across scales means that the degree of guidance is comparable on both submicron and cellular scales.

**Figure 5.**
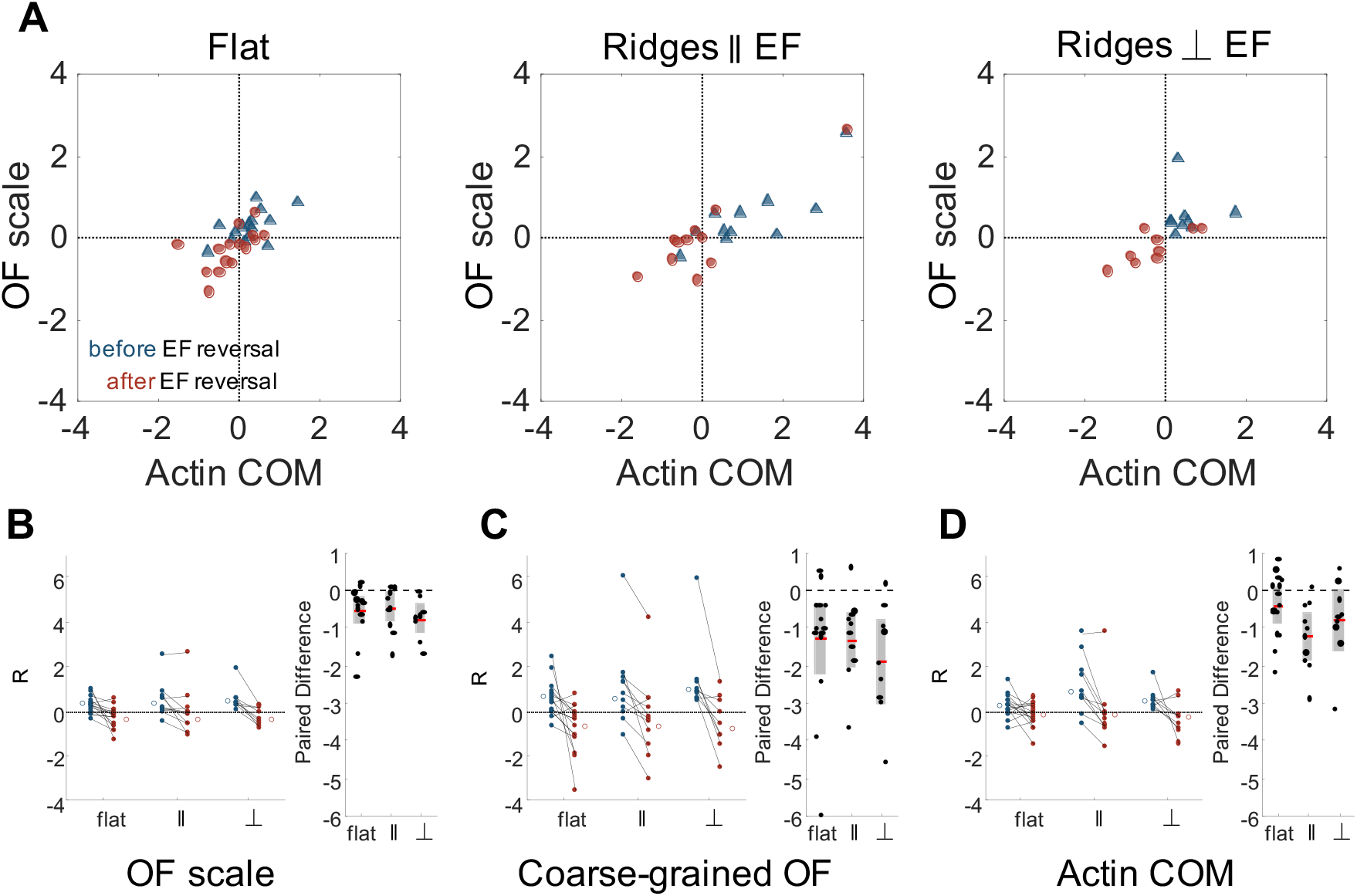
Actin dynamics on multiple scales is correlated, and responds to changing EF direction, independent of nanotopographic guidance. To compare EF guidance of actin waves and, therefore, the cell polarity across substrate type and analysis method, the angular distributions were split by angle, and the logarithm of the ratio of leftward angle counts to rightward angle counts (*R*) was calculated. Positive *R* indicates flow towards the left and negative *R* indicates flow towards the right. (A) Comparison between the small-scale optical-flow measurement (Fig. 4A,B) and the whole-cell, actin-center-of-mass measurement (Fig. 4E,F) before (blue) and after (red) EF reversal. Combined results of cell-scale polarity before and after EF reversal at (B) optical-flow scale, (C) the coarse-grained OF scale, (D) the cellular scale, and their paired differences, respectively.

Because cells exhibit a wide range of guidance ratios, we further assessed the effect of the EF reversal on a cell-by-cell basis. Figs. 5*B-D* show the change in *R* for the three length scales, and for the three experimental conditions. Additional analysis of the paired difference shows that both at the OF scale (Fig. 5*B*) and coarse-grained scale (Fig. 5*C*), the actin dynamics follows the direction of the EF. The cell-scale actin dynamics (COM) (Fig. 5*D*), for which we have fewer data points due to the nature of the measurements, show a significant switch only on ridges parallel to the EF.

## Discussion

We have investigated actin polymerization in the presence of two distinct guidance cues, nanotopography and EFs. Actin dynamics was quantified through an optical-flow-based technique, which allows for determination of preferred directions of actin polymerization on multiple length scales. Our results show that in an EF, actin waves are biased toward the cathode, with stronger guidance when the EF direction is switched than for the initial appearance of an EF. A multiscale analysis (Fig. 3-4) reveals that the preferred direction of actin waves is along the direction of nanoridges, but that the preferred direction is biased by the EF toward the cathode even when the nanoridges are perpendicular to the EF direction. Although cells migrate preferentially toward the cathode on average, we also found that some cells move preferentially toward the anode. In the latter cells, the actin dynamics is also directed toward the anode, consistent with the motion of the cell (Fig. 5). Because no other guidance cues are imposed, our observation of “wrong-way” dynamics of actin and cells point to the persistence of cell polarity.

This coupling of dynamics across scales can be understood conceptually from Fig. 1, which shows that actin polymerization occurs predominantly at the edge of the cell, driving local protrusions and motility. On flat substrates, a single actin wave typically covers the entire leading edge, resulting in whole-cell polarization and migration in electrotaxis; similar behavior has also been observed in chemotaxis (32). Our observations on actin waves on flat substrates reveal three typical cell responses to an EF reversal: (1) unpolarized cells generate new actin polymerization regions toward the cathode; (2) polarized cells can make a U-turn while remaining polarized; and (3) polarized cells can also become rounded and stop, in a so-called “cringe response,” followed by migration in the direction of the EF (33–35).

Nanotopography triggers nucleation and propagation of one-dimensional actin waves (26). This process of esotaxis enables multiple, 1D waves to exist within one cell, as seen in Figs. 1 and 2. An EF can guide these waves but does not alter the 1D wave character. When the EF is aligned parallel to the nanoridges, esotaxis enhances EF guidance of actin dynamics. When the EF is perpendicular to the nanoridges, 1D waves are guided by nanoridges, but the location of new actin waves is biased towards the cathode, leading to overall guidance by the EF. These results demonstrate that actin nucleation, and thus the preferred direction of cell motion, is guided by EF, and that this guidance can be distinct from the local guidance direction of actin waves provided by nanoridges.

Our experiments on EF reversal assess the ability of actin waves to change direction. Because actin polymerization occurs via excitable waves that have a refractory period, individual waves cannot reverse direction. On flat substrates the waves can execute broad turns. In contrast, on nanoridges, the 1D waves cannot turn. The response to a change in the direction of the EF therefore involves new actin waves at locations distinct from the original wave front. On a cellular scale, these multiple new actin waves can recreate the familiar U-turn phenotype.

Our work reveals a mechanism through which actin waves can integrate multiple physical guidance cues. Nanotopography acts on a submicron scale, resulting in 1D waves that are guided along ridges. EFs act on the cellular scale to bias the direction of actin nucleation, and therefore cell migration, toward the cathode. Neutrophils may harness this integration of multiple cues *in vivo* in wound-healing contexts. These cells can navigate both the local microenvironment by following collagen fibers over short distances, and at the same time maintain overall guidance toward wound based on cues that operate on a longer length scale, such as EFs. Because the dynamic cytoskeleton drives a wide range of cell behaviors, the dissection of the response of actin waves to different guidance mechanisms across multiple time and distance scales is a promising approach for understanding the integration of a wide range of guidance cues. Such insights may enable targeted combinations of local and global guidance that could pave the way for flexible means of controlling a wide range of cell behaviors.

## Materials and Methods

HL-60 YFP-actin cells were a gift from the lab of Dr. Orion Weiner of the University of California, San Francisco. The cells were cultured in RPMI 1640 medium, Glutamax (Life Technologies) supplemented with 10% heat-inactivated fetal bovine serum (Gemini Bio). The cells were kept in a humidified atmosphere at 37 °C and 5% CO_2_. Cells were differentiated to be neutrophil-like with a supplement of 1.3% dimethyl sulfoxide Hybri-Max (Sigma Aldrich) at 4.5 × 10^5^ cells/mL.

To create the nanotopography, we used multiphoton absorption polymerization (36). Molding techniques were employed to create many acrylic polymer replicas of the nanotopographic substrates (37). Surfaces were coated with 10 μg/mL fibronectin (Sigma Aldrich) for 1 hour.

Cells were resuspended in modified Hank’s Balanced solution (mHBSS, Sigma-Aldrich) with 1 µM N-Formyl-Met-Leu-Phe (fMLP, Sigma Aldrich), and then were then plated in a 3D-printed electrotaxis chamber composed of polylactic acid (38). Toxicity and changes in pH due to electrode byproducts were minimized by using separate baths for each AgCl electrode and the cell-mHBSS solution, and by connecting the baths using agar bridges in mHBSS (2%, 0.5 cm diameter, created with glass tubing). EF polarity switches were applied over a period of < 10 sec. Time-lapse imaging was performed using a Perkin Elmer spinning-disk confocal microscope.

Our analysis of actin-wave dynamics was based on an optical-flow algorithm, as described previously (25). Briefly, this technique allows the flow of actin to be quantified in an unbiased manner. Optical flow uses the intensity of actin fluorescence images to calculate spatial and temporal gradients and to determine the ground-truth intensity flow. We used the Lucas-Kanade approach with a varied weight matrix, a 4 μm × 4 μm Gaussian with a standard deviation of 0.63 μm for small-scale flow and a 12.8 μm × 12.8 μm Gaussian with a standard deviation of 2.1 μm for larger-scale flow (31).

To study the changing shape of migrating cells, we applied a snake (active-contour) algorithm, as described previously (29). The snake algorithm was used to extract the boundary (with 200 boundary indices) of the binarized cell with a 1:1 mapping between adjacent frames, so that one index position (50) was always oriented directly leftward. The location of the cell centroid and averaged actin intensity for each boundary index was calculated through time.

## Acknowledgments

We thank the University of Maryland Imaging Incubator Core Facility for use of their systems in collecting images for this work. This work was supported by the Air Force Office of Scientific Research MURI grant FA9550-16-1-0052. We thank the MURI team for fruitful discussions. A.B. and L.C. were supported by COMBINE NRT award number 1632976. M.J.H was supported by NSF award number 1806903. We thank Dr. Orion Weiner at the University of California, San Francisco for gifting us the HL60 cell line.

