## Supplemental Information for "Actin dynamics as a multiscale integrator of cellular guidance cues"

**Supplementary Information for**
### Actin dynamics as a multiscale integrator of cellular guidance cues

Abby L. Bull[1,2], Leonard Campanello[1,2], Matt J. Hourwitz[3], Qixin Yang[1,2], Min Zhao[4], John T. Fourkas[1,3], Wolfgang Losert[*1,2]

[1]Institute for Physical Science and Technology, University of Maryland, College Park, MD
[2]Department of Physics, University of Maryland, College Park, MD
[3]Department of Chemistry and Biochemistry, University of Maryland, College Park, MD
[4]Institute for Regenerative Cures, Department of Ophthalmology & Vision Science, Departments of Dermatology, School of Medicine, University of California, Davis, CA, 95616

*Corresponding Author: Wolfgang Losert


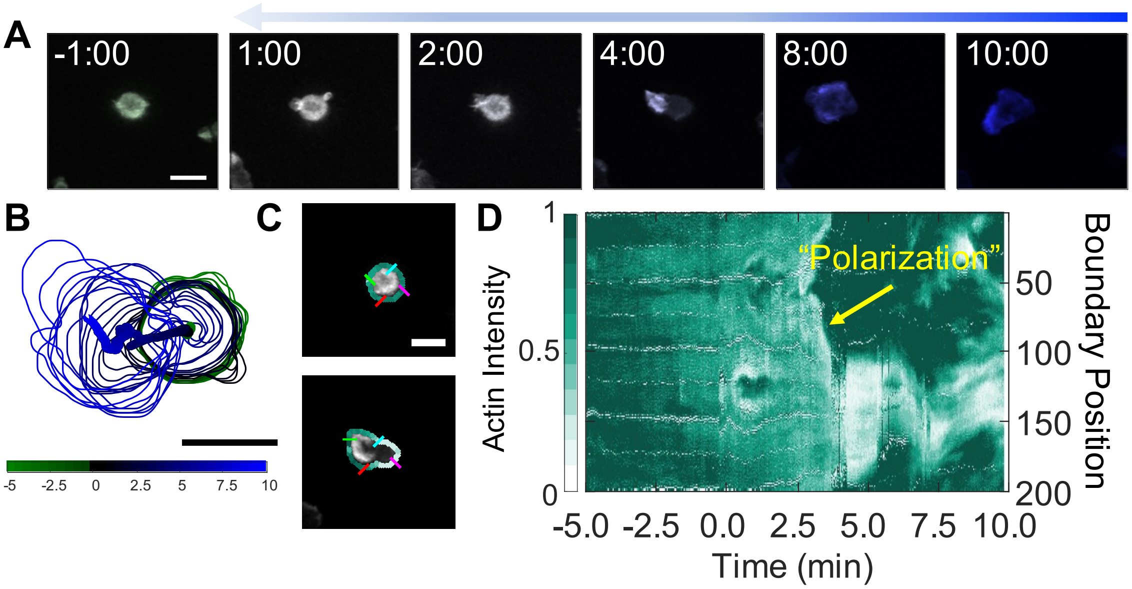


**Fig. S1.** Activation of an HL60 cell by introduction of an EF. (A) Time-lapse snapshots of differentiated YFP-actin-labelled HL60 cell on a flat substrate. There is no EF present at negative time. The cathodal direction is to the left in positive time. The scale bars are 10 µm. (B) The boundary shape and path of the centroid of the cell from (A) from -5:00 min to 10:00 min in 0.5-min increments, with (C) an example of the extracted shape of the cell colored by actin intensity. (D) The evolution of the normalized boundary actin fluorescence intensity visualized as a kymograph. Indicated with the yellow arrow is the general event of polarization of the cell. From a rounded shape with generally uniform fluorescence to an elongated shape with extending actin-rich lamellipodia, the cell becomes activated (i.e., has a polarization event) between 2.5 and 5.0 min. Many boundary regions of the cell become darker green after this event (at boundary positions 0 to 100) indicating the increase in actin intensity in the lamellipodia while other regions (from 100 to 200) decrease in actin intensity as they become the back of the cell after symmetry breaking.

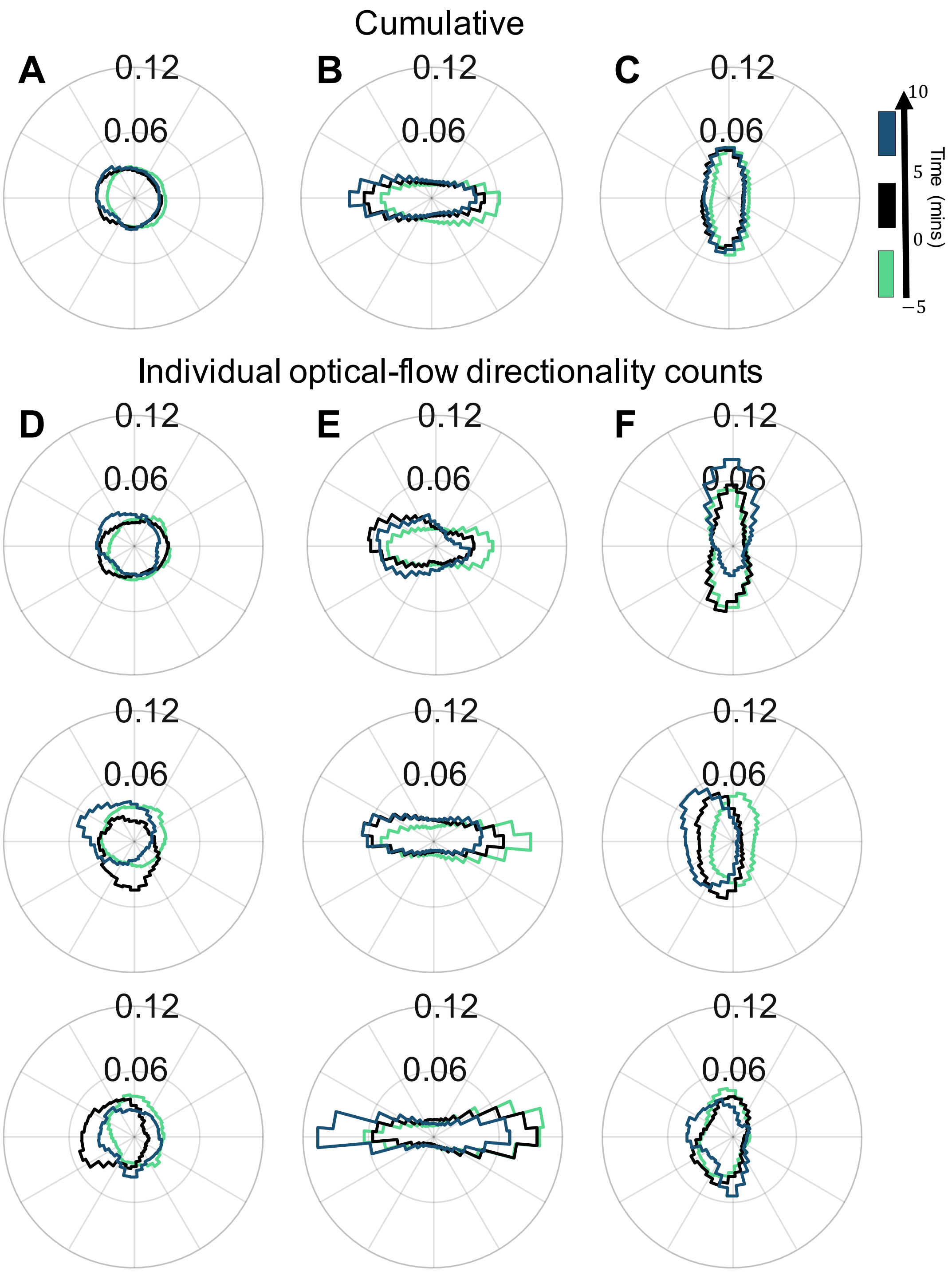


**Fig. S2.** Analysis of actin dynamics analyzed before and after the introduction of an EF. (A) Cumulative flow-direction distribution shown for the time ranges (green) 5 min before EF introduction; (black) 5 min after EF was switched on; and (blue) the period 5 min to 10 min after the EF was switched on averaged for all cells on (left) flat substrate, (middle) nanoridges parallel to the EF, and (right) nanoridges perpendicular to the EF. Three characteristic examples of individual, cell-normalized distributions (in counts) of optical-flow directions on (D) a flat substrate, (E) nanoridges parallel to the EF and (F) nanoridges perpendicular to the EF.
